# Altered Cardiac Neural Crest Migration Patterning in a Left Atrial Ligation Model of Hypoplastic Left Heart Syndrome

**DOI:** 10.64898/2026.03.11.711140

**Authors:** Aining Fan, Robert Porter, Hyun Maeng, Stephanie E. Lindsey

## Abstract

Cardiac neural crest cells (CNCCs) contribute to key cardiac structures during embryonic development. Disruption of CNCC patterning or function can lead to congenital heart defects. Here, we investigate whether hemodynamic perturbation alters CNCC behavior in chick embryos. We use the left atrial ligation model to modify intracardiac blood flow in the early common-atrium, common-ventricle heart and track retrovirally labelled CNCCs for lineage tracing and single-cell transcriptomic analysis. Results revealed a significant reduction of CNCC derivatives in major cardiac regions, including the pharyngeal arch arteries and myocardium, in flow-perturbed embryos compared with controls. Notably, despite reduced CNCC numbers in the PAAs, their relative proportion increased, suggesting retention within the PAAs and delayed differentiation. Transcriptional analysis shows the expression of CNCC post-migratory markers (HAND1, FOXC2, GATA6, and TBX2) were consistently downregulated at 4, 24, and 48 hours after LAL. Together, these findings indicate that hemodynamic perturbation impairs CNCC migration and differentiation while preserving their capacity to contribute to mature cardiac structures.

## 1 INTRODUCTION

Cardiac neural crest cells (CNCCs) are integral to normal cardiac development, initiating cardiac septation and ultimately forming the tunica media of the great vessels ^1^. The crest cells receive fate instructions from their origin as well as respond to their local environment ^2,3^.Deviations in CNCC migration patterns or cellular function can lead to a number of cardiac abnormalities ^4,1,5,6^. Cells of the cardiac neural crest migrate out of the caudal neural tube (somites 1-3), pause in the circumpharyngeal ridge and seed the arch arteries to make up their ectomesenchyme ^7,8,9^. A subset of CNCCs, originating between the fourth and sixth arch arteries, extend dorsally into the aortic sac and initiate the formation of the aortic pulmonary septum ^10^. The shelf spirals into the distal outflow tract setting the prevalvular cushions and ventricle up for septation. ^11^. Neural crest cells remaining in the arch arteries form the smooth muscle tunics of the pharyngeal arch artery derivatives and the distal continuations of the great arteries.

Crest ablations are associated with failure of arch arteries III, IV (right), and VI to develop to the proper size ^12^ and loss of pharyngeal arch artery bilateral symmetry ^13^. Ablation of a pre-mirgratory CNCCs results in a high incidence of persistent truncus arteriosus and ventricular septal defect ^6^, while double outlet right ventricle with ventricular septal defect results from small lesions of cardiac neural crest ^14^. CNCCs are not required for the formation of the arch arteries, but have been shown to be critical to their maintenance ^13^. Neural crest ablation has also been shown to influence the cells of the secondary heart field, which contribute to the myocardialization of the outflow tract ^1^ as well as contribute to cardiac myocardium ^15^. In crest-ablated embryos, the cells that normally migrate from the secondary heart field proliferate rather than migrate and differentiate into myocardium ^1^. The fate of the pharyngeal arch arteries (PAAs) and the CNCCs are interrelated. The avian arch arteries are right-side dominant, with the right fourth PAA forming part of the aortic arch, the left fourth PAA forming the aortic arch in mammals. Jiang et al, discovered a right-sided aortic arch mouse embryo, which involves the retention of the right fourth arch artery as the arch of the aorta, the right sixth arch artery as the ductus arteriosus, and the right dorsal aorta as the descending aorta, with a neural crest pattern that resembled that of the left arch arteries ^16^. Solovieva & Bronner found that ablation of right cardiac neural crest resulted in embryos with missing arteries ^3^.

Changes in geometry, following neural crest cell ablation, also lead to changes in flow and cardiac function. Embryos were characterized by decreased systolic and diastolic blood pressures following ablation of the entire length of CNCCs ^17^, as well as depressed shortening and ejection fractions following ablation of CNCCs destined for arch III and IV ^18^. Embryos where targeted crest ablation was performed on cells destined for arch III and IV exhibited decreased blood flow or absence of blood flow in the right 4th PAA, altered conotruncal shape, depressed contractility and dilation of the primitive ventricle, decreased emptying of the bulbus cordis, incompetent truncal cushions and incomplete looping of the cardiac tube ^19^.

In the present study, we examine the effects of changing the hemodynamic environment, specifically inducing a single ventricle formation through the classical chick left-atrial ablation model, ^20^ on CNCC migration, patterning and differentiation. With the left atrial ligation (LAL), intracardiac flow patterns from the right cardinal vein, right vitelline vein and left vitelline vein are altered immediately, leading to reduced wall shear stress at the left atrioventricular canal and left side of the common ventricle ^21^. The redistributed intracardiac blood flow adversely alters aortic arch artery development ^22^, leading to aortic stenosis and malpatterning. We use the LAL model and retrovirally labeled CNCCs to test the hypothesis that redistributed intracardiac blood flow alters CNCC migration patterning.

## 2 RESULTS

### 2.1 Left atrial ligation alters cardiac anatomy and neural crest cell migration patterns

To assess how CNCCs adapt to altered mechanical signaling, we first labeled CNCCs with a replication incompetent avian retrovirus while the cells were confined to the neural tube (HH8-HH10) ^23,15^ (Figure 1A). We then performed a LAL surgery at HH21 in which a suture was tied around the left atrium and pulled tightly enough to arrest flow ^20^ (Figure 1B). The LAL model was chosen because of its well documented hypoplastic left heart phenotype and mechanical characterization ^24,25,26,27^. Whole embryo stereoscope and tissue section analysis was conducted to confirm LAL defect (***Figure 2***). Large structural defects exist within 72 hours post-LAL surgery (***Figure 2***)B,C. Light-sheet fluorescence microscopy confirmed pronounced four-chamber structural differences between control and LAL embryos (***Figure 2C***), with LAL embryos exhibiting a reduced left ventricular lumen accompanied by left ventricular wall thickening. These morphological features resemble hypoplastic left heart syndrome (HLHS), consistent with the well studied LAL model ^20,28,24^. The replication-incompetent avian retrovirus (RIA) enabled stable, heritable labeling of host cells and their progeny at single-cell resolution ^23^. To confirm viral labeling, embryos injected with membrane-targeted YFP retrovirus were dissociated at 48 hours post-injection. Single cells were plated and imaged individually, revealing robust membrane-localized green fluorescence, confirming successful viral infection (***Figure 6***). Following validation, virally injected embryos were allowed to develop until embryonic day 6 (HH29), after which they were harvested, cryosectioned, and imaged by fluorescent microscopy.

**FIGURE 1.**
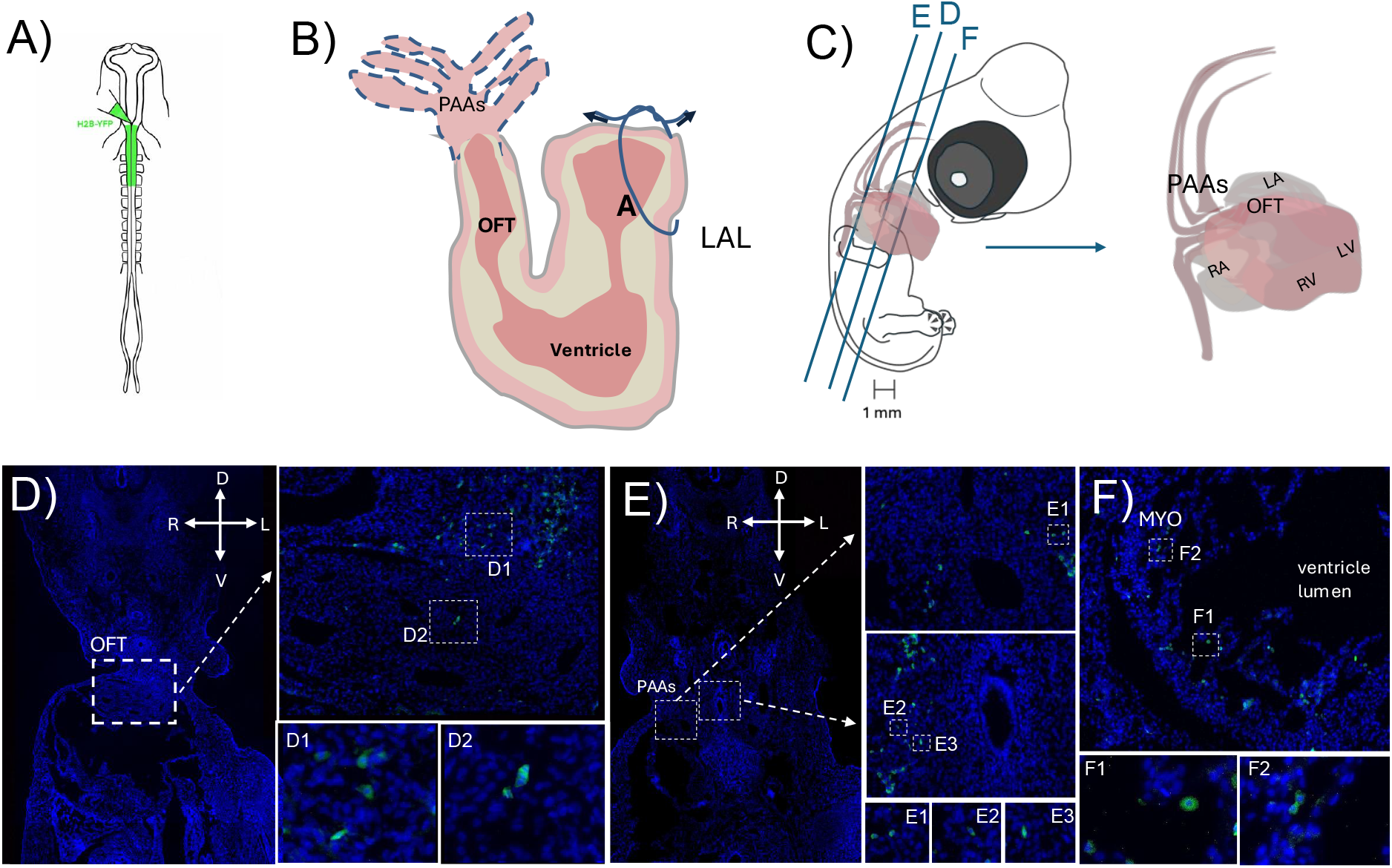
Retroviral labeling of CNCCs during early cardiogenesis. **(A)** Schematic of experimental workflow: Replication-incompetent avian (RIA) retrovirus encoding membrane-targeted YFP injection into the neural tube at the level of the anterior three somites beneath the hindbrain. **(B)** Left atrial ligation surgery performed at HH21 (day 3.5) **(C)** HH29 embryo (day 6) with guides for histological section planes and enlarged view of heart. Blue lines indicate the planes corresponding to the sections shown in (D–F). **(D-F)** Transverse sections highlighting regions of interest, including the outflow tract (OFT), pharyngeal arch arteries (PAA), and myocardium (MYO). D, dorsal; V, ventral; L, left; R, right. Dashed boxes denote the relative locations of membrane-YFP labeled cell populations. **(D1-2, E1-3, F1-2)** Higher-magnification views of the boxed regions in (D–F), respectively.

**FIGURE 2.**
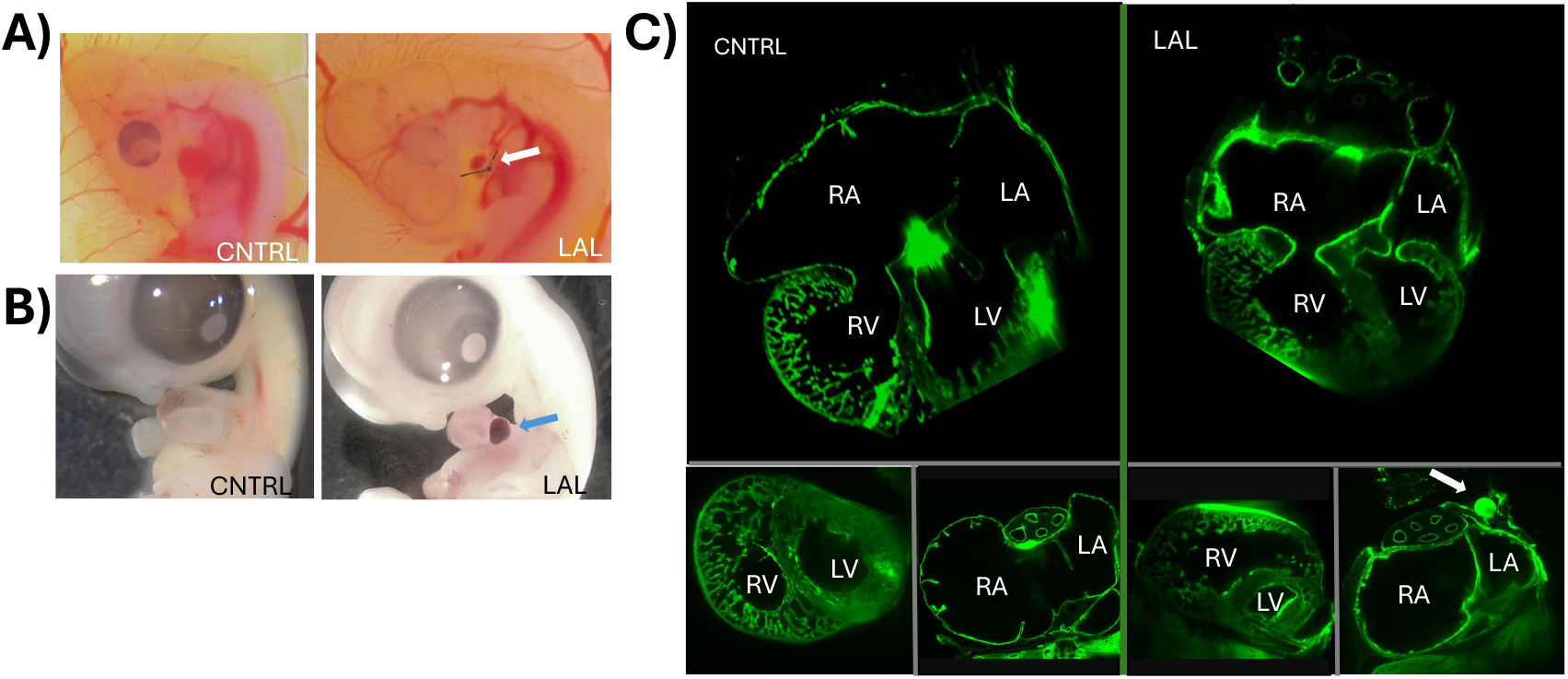
Representative control and LAL embryos at (a) HH24 (day 4) and (b) HH30 (day 7) c) Four chamber heart section views (top) and short-axis views of ventricles and atria (bottom) in control and LAL HH30 hearts. Arrows(white, blue) highlight the knot tying off the left atrium. RA-right atrium, LA-left atrium, RV-right ventricle, LV -left ventricle, CNTRL - control, LAL-left atrial ligation

In keeping with previous studies ^13,15^, viral-labeled neural crest–derived cells were observed within key cardiac structures, including the pharyngeal arch arteries, outflow tract, and ventricle myocardium (***Figure 1 D–F***). Individual structures were further resolved at higher magnification, where the virus-labeled cell nuclei were identified by DAPI staining, while the cell membranes were delineated by green fluorescence (***Figure 1D1–2, E1–3, F1–2***). Similar labeling patterns were observed in embryos injected with H2B-YFP retrovirus (***Figure S1***). Virus-labeled CNCCs were detected surrounding the OFT (***Figure S2A1***) and within the myocardium, but were notably absent from the epicardium (***Figure S2A2***).

To assess the effect of LAL on CNCC migration and differentiation, HH29 LAL embryos and stage-matched controls were processed for histological analysis. We stained for known CNCC derivatives, smooth muscle actin for the tunica media of the PAAs, troponin-T and myosin heavy chain for the ventricle. YFP-labeled CNCCs were identified within the PAA tunica media along with smooth muscle action in both control and LAL embryos (***Figure 3A-D***), indicating that CNCCs continues to differentiate in smooth muscles cells post LAL. YFP-labeled CNCC derivatives were also detected in the ventricular myocardium of LAL embryos, co-localized with troponin-T and myosin heavy chain (***Figure 1 E***), consistent with previous studies ^15^. These observations indicate that the migratory path and differentiation patterns of CNCCs is largely preserved following LAL -induced hemodynamic perturbation. However, quantitative analysis revealed a sustained reduction in CNCC abundance within these anatomical regions across multiple developmental stages following LAL intervention (***Supplementary Table 1***). While reduction in CNCC cell count was seen across stages, statistical significance was only found for HH29 PAAs and ventricular myocardium regions (***Figure 4 C***). Results indicate a delay in CNCC migration and differentiation progression while maintaining the same capacity to contribute to mature cardiac structures.

**FIGURE 3.**
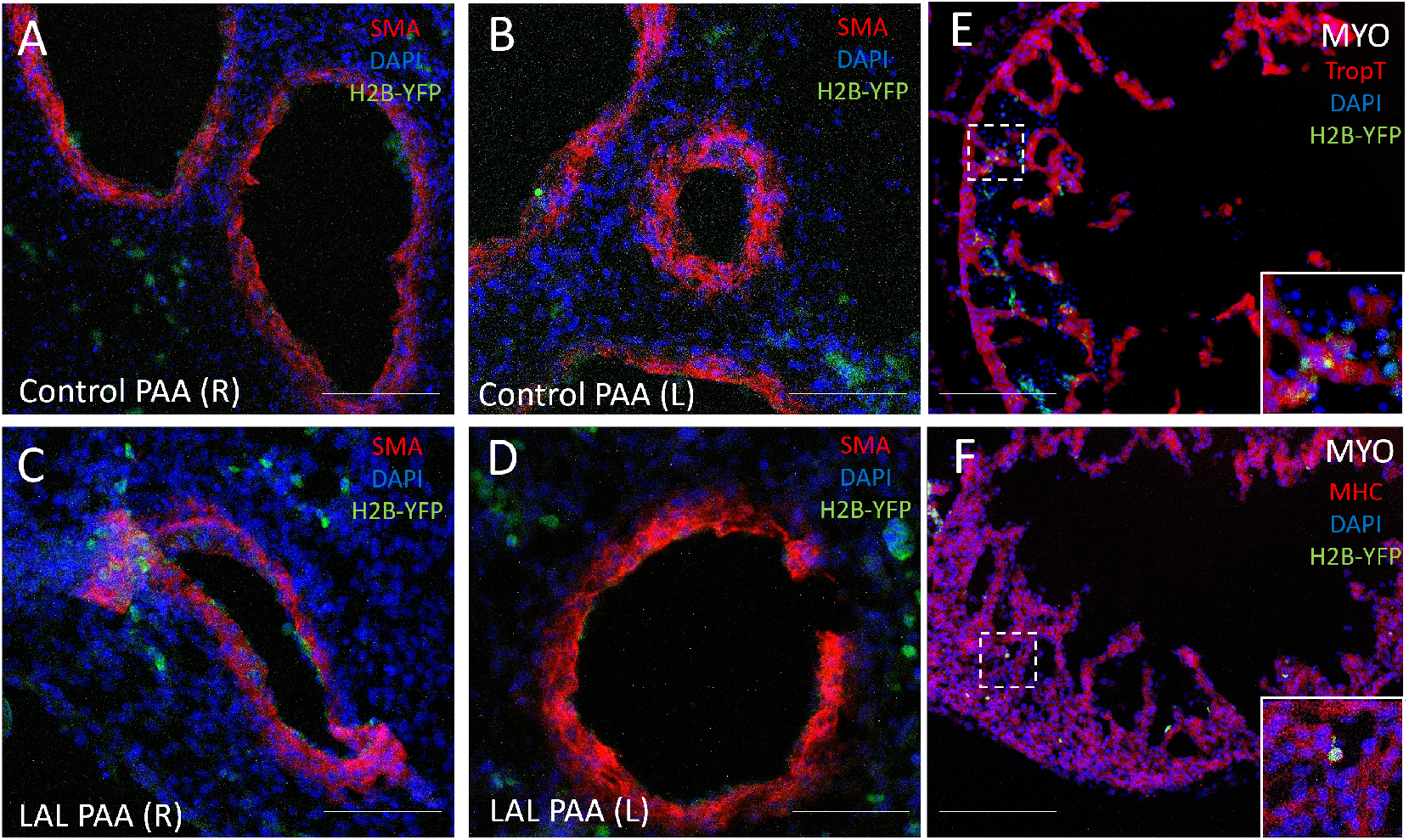
Comparison of smooth muscle differentiation of CNCCs in flow-perturbed and control chick embryos. **(A–D)** Retrovirally labeled CNCC-derived cells expressing H2B–YFP (green) were identified within the PAAs and assessed for expression of the smooth muscle marker smooth muscle actin in HH29 embryos. **(A)** Right PAA in a stage-matched control embryo. **(B)** Left PAA in a stage-matched control embryo. **(C)** Right PAA in a left atrial ligation (LAL) embryo. **(D)** Left PAA in an LAL embryo. **(E, F)** CNCC derivatives were also observed in the ventricular myocardium of HH26 LAL embryos and were co-stained with cardiac troponin T **(E)** and Myosin Heavy Chain **(F)** in LAL embryo at HH26. Scale bar: 100*µ*m; Image scale:A-F: 436 *µ*m × 366 *µ*m

**FIGURE 4.**
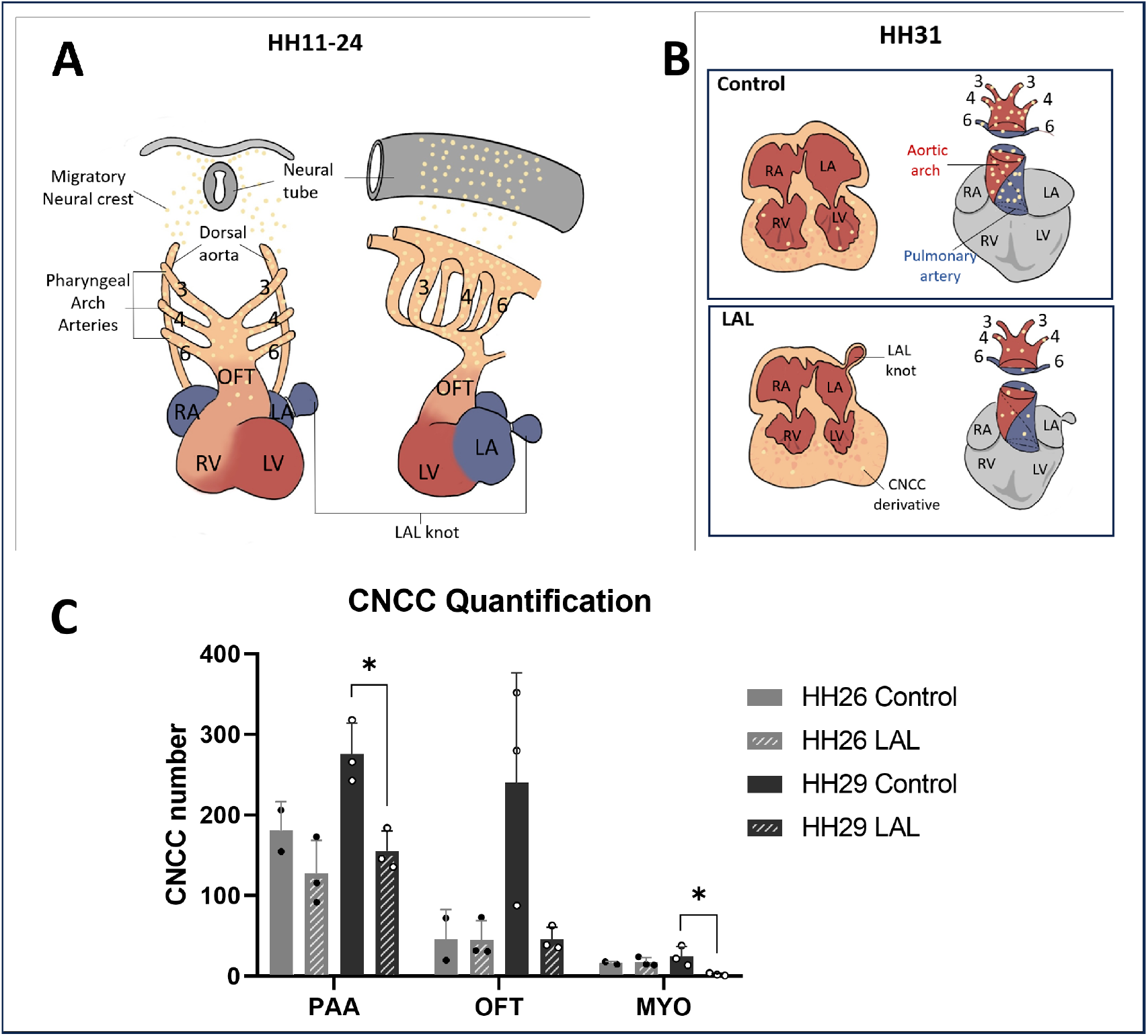
CNCC enrichment and distribution in the PAA, OFT, and myocardium are reduced in LAL embryos. **(A)** Schematic illustration of control CNCC migration from HH11 to HH24. CNCCs originate from the dorsal neural tube and migrate through PAA 3, 4, and 6 before a subset of cells initiates cadiac septation in the outflow tract. LAL was performed at HH24 by placing a suture around the left atrium to restrict blood flow. **(B)**Schematic representation of reduced CNCC derivatives in the OFT and ventricular myocardium at HH31 in LAL embryos compared with stage-matched controls. **(C)** Quantification of CNCCs in the PAA, OFT, and myocardium MYO at HH26 and HH29. CNCC numbers were counted from three consecutive sections per embryo and averaged. Sample size: n = 3 embryos for HH26 LAL, HH29 control, and HH29 LAL; n = 2 embryos for HH26 control. Statistical significance was assessed using an unpaired t-test with Welch’s correction (p < 0.05). Abbreviations: LV, left ventricle; RV, right ventricle; LA, left atrium; RA, right atrium; OFT, outflow tract; PAA, pharyngeal arch arteries; MYO, myocardium.

### 2.2 Transcriptomic alterations in hemodynamically perturbed hearts revealed by single-cell RNA sequencing

To capture the early transcriptional response of CNCCs to hemodynamic perturbation, we compared virally labeled CNCC populations between LAL embryos and stage-matched controls at HH24 (day 4), roughly 4 hours post-LAL. Differential gene expression analysis was performed on log-normalized single-cell expression values, and significant genes were identified based on adjusted statistical thresholds (***Figure 5 A***). The volcano plot revealed a subset of genes significantly downregulated in LAL CNCCs relative to controls, including multiple canonical regulated genes associated with neural crest migration and early differentiation. Notably, key migratory transcription factors such as SNAI1, ETS1, TAF10, TFAP2A, FOXC2, HAND1, HAND2, and MAFB were among the downregulated genes (***Figure 5 A, Supplementary Table 2***) ^29 30 31^, suggesting that flow perturbation alters transcriptional programs involved in CNCC developmental progression.

**FIGURE 5.**
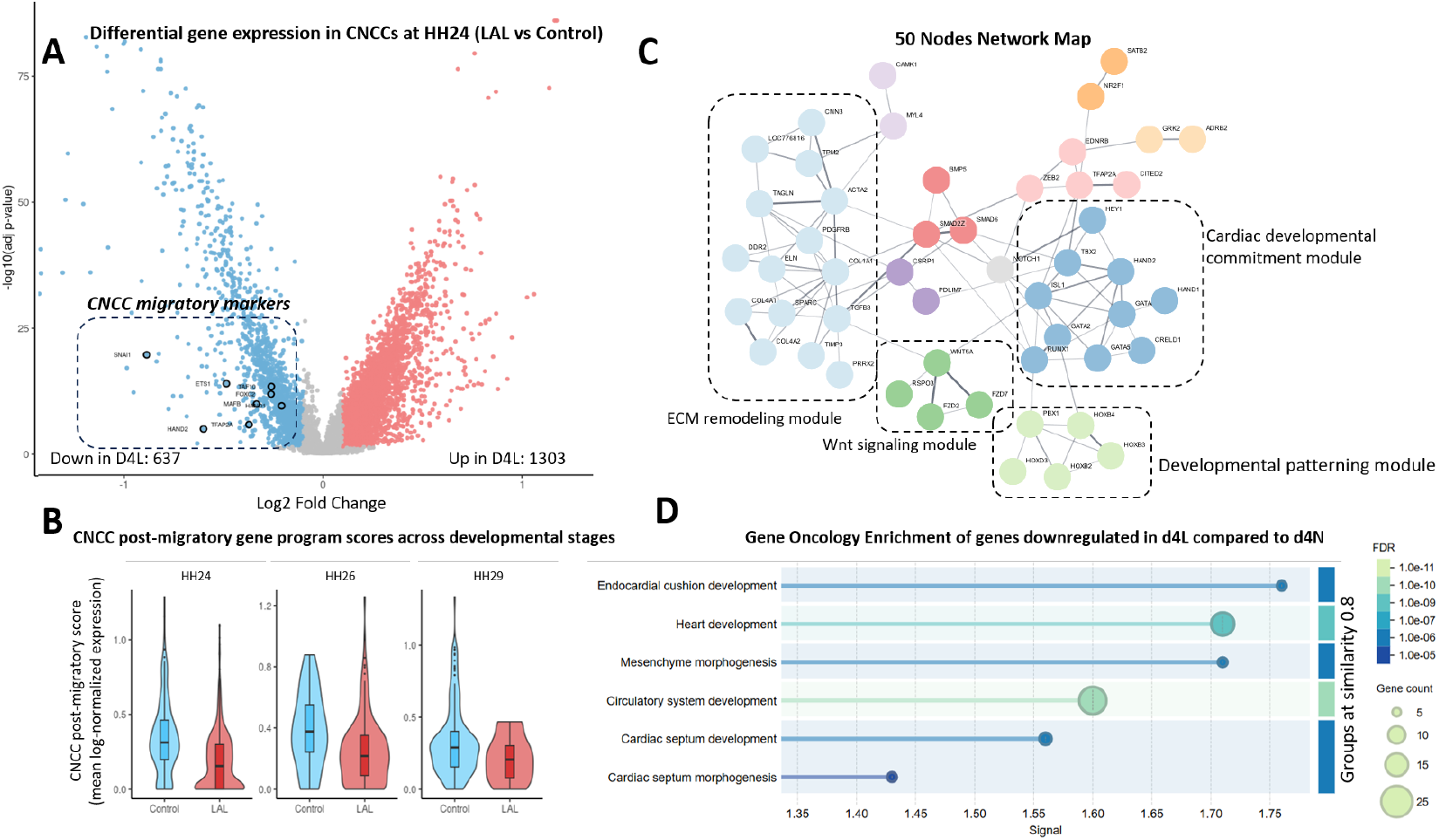
Transcriptional maturation of CNCCs. (**A**) Volcano plot showing differential gene expression between HH24 (day 4) LAL and stage-matched control CNCCs based on single-cell RNA sequencing of FACS-sorted Tg(H2B-RFP)+ cells. Selected canonical CNCC migratory regulators are highlighted. (**B**) Distribution of post-migratory CNCC gene program scores across developmental stages from HH24 (day 4) to HH29 (day 6). Protein–protein interaction network constructed from 50 genes selected from the downregulated differentially regulated genes list. Markov clustering (MCL) was used to identify network modules.(**D**) Gene Ontology (GO) biological process enrichment analysis of genes downregulated in HH24 LAL CNCCs.

To determine whether these transcriptional changes correspond to altered maturation dynamics of CNCC, we quantified a post-migratory CNCC gene program score for each cell, defined as the mean log-normalized expression of HAND1, FOXC2, GATA6 and TBX2. Across developmental stages from HH24-HH29 (day 4 to day 6), CNCCs from LAL embryos consistently exhibited lower post-migratory program scores compared with control embryos, with the distributions shifted toward reduced expression levels at each stage (***Figure 5 B***). These results indicate a persistent attenuation of post-migratory differentiation programs in CNCCs following LAL treatment.

To further examine whether the downregulated genes form coordinated regulatory networks, we constructed a protein-protein interaction (PPI) network and visualized the resulting network. 50 genes selected from the downregulated differentially expressed genes list formed a highly interconnected network consisting of 50 nodes and 101 edges, significantly exceeding the 8 edges expected by chance (PPI enrichment *p* < 1.0 × 10^−16^), indicating a strong functional connectivity. Network clustering further resolved multiple subclusters, with four major functional modules corresponding to extracellular matrix organization, cardiogenic differentiation, embryonic pattering, and Wnt signaling regulation (***Figure 5 C***).

Gene oncology further revealed significant over-representation of cardiovascular developmental processes, including endocardial cushion development, mesenchyme morphogenesis, circulatory system development and cardiac septum development (***Figure 5 D***). Collectively, differential expression and PPI network analyses confirm disruption of cardiogenic regulatory modules, while the reduced activation of post-migratory CNCC gene programs across developmental stages further suggests that altered blood flow delays CNCC maturation rather than fundamentally changes their lineage identity.

## 3 DISCUSSION

### 3.1 Hemodynamic forces and CNCC behavior in cardiac morphogenesis

The etiologies of congenital heart defects remain poorly understood ^32,33^, but are believed to result in part from hemodynamics ^34,35^. Hemodynamic forces play a critical role in cardiac morphogenesis, whereby biomechanical cues generated by blood flow influence cellular differentiation, migration, and tissue patterning ^36,37,38,39^. In the present study, we use the chick left atrial ligation model of hypoplastic left heart syndrome to study the effects of the changing biomechanical landscape on CNCCs. HLHS is associated with a hypoplastic left ventricle, mitral valve hypoplasia or stenosis, aortic atresia, and hypoplasia or coarctation of the aorta ^40^. These phenotypes are reproduced in the chick LAL model where altered ventricular filling in early development results in altered ventricular function and morphology, along with great vessel defects ^20,28,22^. In the clinical patient, retrograde aortic flow is thought to be responsible for impaired development of the aortic root and ascending aorta ^41^. Ocillatory flow with reduced flow velocity has also been documented in the HH28 (day 5.5) LAL ventricle (^24^), and may be responsible for the changes seen between HH26 and HH29 LAL CNCC count when compared with controls. Retrograde flow is associated with other great vessel malformations ^42^ and alters gene expression, as directional changes in wall shear stress over the cardiac cycle are sensed by endothelial cells ^43^.

CNCCs are essential to cardiovascular development ^44,15^. Neural crest cell ablation studies have produced the same range of defects that arise from mechanical perturbation studies ^45,12,46^, supporting a potential interplay between CNCC behavior and mechanical cues. Following ablation of premigratory neural crest destined for PAA III, PAA IV and the OFT septum, altered arch artery, pharyngeal mesenchyme, cardiac looping and OFT septation defects arise ^46^. CNCC ablations have also led to loss of bilateral symmetry, with some arches appearing small or occluded, reduced quantity of pharyngeal mesenchyme, and hypoplasia of the fourth arch artery ^12,17^. Similar defects develop from HH18 arch artery occlusion, vitelline vein ligation, or left atrial ligation ^47,38,20^. With neural crest ablation, functional changes in hemodynamics precede abnormal structural changes in the heart and aortic arch arteries ^17^. Indeed, genetic models of cardiac malformations perturb hemodynamics which contribute to the severity of the defect^48,19,50,4^. Structure and function are integrally related in the cardiovascular system. Although both hemodynamic perturbation and CNCC ablation independently recapitulate phenotypic features of congenital heart disease, their mechanistic interdependence has not been systematically investigated.

In the present study, we demonstrate that hemodynamic perturbation directly impacts CNCC migratory progression and subsequent lineage contribution. Using retroviral lineage tracing and immunohistochemistry, we confirmed the spatial distribution of CNCC derivatives within the pharyngeal arch arteries, outflow tract, and myocardium in wild-type chick embryos as previously reported ^15^ (Figure 1). While CNCCs follow the same migratory path through the PAAs, OFT and myocardium, the number of cells in each region is greatly reduced. A higher percentage of cells (76%) remain the the PAA region in HH29 LAL embryos compared to (53%) in controls. Roughly 22% of CNCCs are present in the OFT in HH29 LAL embryos, as the heart completes septation, compared to 42% of controls (Figure 4, Table S1). Transcriptomic analysis revealed significant downregulation of canonical CNCC regulatory and migratory genes (HAND1, GATA5/6, MAFB, FOXC2, TBX2) shortly after LAL intervention. Moreover, post-migratory gene expression in LAL embryos remained progressively downregulated across developmental stages, indicating sustained impairment of CNCC migratory progression. This persistent suppression ultimately corresponds with the reduced CNCC distribution in key cardiac structures observed in our quantitative analyses.

### 3.2 Mechanotransduction links flow peturbation to CNCC dysfunction

Shear stress and shear stress -activated or repressed gene expression are important factors in remodeling of the cardiovascular system ^51,52^. Though many markers of CNCCs in different states (migratory, premigatory, post-migratory) have been identified ^29^, the role of hemodynamic signaling in CNCC migration and differentiation is poorly understood. KLF2 is a shear stress–responsive transcription factor expressed in endothelial cells of the developing heart and serves as a key mediator linking hemodynamic forces to transcriptional regulation ^50,53,54^.

KLF2 directly regulates several cardiac developmental genes, including SOX9, GATA4, and TBX5, and its deficiency leads to vascular and valvular defects ^55^. HEY2 is a downstream effector of the Notch signaling pathway, which functions as a mechanotransducer during cardiac development ^56,57,58^. In zebrafish models, reduced contractile force and altered wall shear stress have been shown to suppress Notch1b activity, resulting in ventricular valve malformations ^59^.

In our dataset, both KLF2 and HEY2 exhibited reduced expression in LAL embryos (***Supplementary Table 2***), suggesting disruption of flow-sensitive signaling pathways under hemodynamic perturbation. Salman et al showed that LAL intervention causes an immediate flow disturbance over atrioventricular (prevalvular) cushions ^28^. A possible mechanistic cascade supported by the current study is the post-LAL reduction in atrioventricular cushion shear stress leads to decreased KLF2 activity, which in turn attenuates Notch pathway signaling and downregulates downstream targets. These include chamber-specific regulatory genes such as HEY2 as well as atrioventricular canal–associated genes such as TBX260. Importantly the transcriptional regulators HEY2 and TBX2 have also been implicated in CNCC migration and differentiation ^61,62^. The coordinated downregulation of HEY2 and TBX2 may therefore contribute to impaired CNCC developmental progression and altered cellular distributions within cardiac structures.

### 3.3 Limitations

A number of red blood cells were captured with FACs sorting, resulting in a much lower H2B-RFP positive cell count post-RNA-sequencing than expected. Further transcriptional analyses were constrained by the relatively low number of CNCCs captured for sequencing, with HH26 control and HH29 LAL groups suffering from particularly low cell counts. The LAL model, though well documented, represents a relatively strong hemodynamic perturbation and tissue perturbation. With the LAL, it is difficult, to isolate the effects of hemodynamics alone and know what aspects of LAL induced malformations led to the altered CNCC patterning. Despite these considerations, the present study shows that CNCC patterning is altered following mechanical intervention. Results support an interaction between hemodynamic cues and CNCC migratory behavior that should be further studied. Future investigations incorporating larger transcriptomic datasets, targeted gain- and loss-of-function approaches, and more refined hemodynamic perturbation strategies will further clarify the mechanogenetic pathways linking flow dynamics to CNCC-mediated cardiovascular morphogenesis.

## 4 EXPERIMENTAL PROCEDURES

### 4.1 Cell culture and virus preparation

Avian retrovirus was produced by transfecting embryonic fibroblast DF1 cells (ATCC, Manassas, VA; CRL-3586) with RIA-H2B-YFP plasmid (#96893) or RIA-H2B-RFP plasmid (#92398) and VSVG shell plasmid. Viral lysate was collected over 4 days with aliquots frozen at the -80C. Lysates were filtered and ultracentrifuged at 75500rcf for 2 hours. The supernatant was dried with aspiration, and the pellet was dissolved in 20–30 ml of DMEM, aliquoted, and stored at -80^°^C until the time of injection.

### 4.2 Embryo culture and preparation

Fertilized eggs obtained from commercial farms (USA) were incubated blunt-side up in a continuous rocking incubator at 37.5^°^C and 60% humidity to reach the desired Hamburger Hamilton (HH) stage ^63^. Retroviral injections to label CNCCs were performed *ex ovo* on windowed HH8-HH10 embryos as previously described ^15^. Briefly, using a glass micro-needle fashioned from pulled capillary tubes (0.75 mm ID), the neural tube lumen was injected with a working viral solution to fill the lumen at the level of the mid-otic vesicle to somite three. The working viral solution consisted of Ringer’s solution, food dye (Spectral Colors, Food Blue 002) and viral stock. Embryos were then sealed with surgical tape and returned to a stationary incubator at 37.5^°^C for up to six days of development. Embryos were harvested at HH24 (day 4) HH26 (day 5), and HH29 (day 6).

### 4.3 Left atrial ligation

Using sterilized forceps on windowed HH24 (day 4) embryos that previously underwent viral injection, the overlying chorionic and allantoic membranes were removed. The embryo was gently lifted and rotated vertically with the forceps so that the left side faced upward. With the use of fine tip forceps, the pericardium above the left atrium was opened and a pre-tied 10-0 nylon suture knots (∼0.5 mm in diameter) was positioned around the left atrium. The suture was tightened to reduce left atrium volume by approximately 75%, thereby restricting blood flow through the left side of the atrioventricular canal. The suture ends were trimmed with micro-scissors. The embryo was then returned to its original position, with the right side facing upward, and returned to the incubator to continue developing until the desired stage.

### 4.4 Tissue dissociation and cell sorting

At the desired stages, embryos were dissected away from the egg and placed in cold Ringer’s solution. The PAAs and heart were separated and washed 3x in dPBS. The dissected heart and outflow tract were dissociated using Accumax (Innovative Cell Technologies, Inc.) and were subsequently filtered, and resuspended in presort flow buffer (BD 563503). Cells were sorted using the BD Aria FACS Fusion and Aria BD FACS II (BD Biosciences).

### 4.5 RNA sequencing and Data Analysis

Following cell sorting, the Chromium Next GEM Single Cell 3′ Kit (10x Genomics, PN-1000268) was used to generate barcoded cDNA libraries. Libraries were sequenced at the UCSD IGM Genomics Center on an Illumina NovaSeq 6000 platform. Raw sequencing reads were processed using the Cell Ranger pipeline (v5.0.1, 10x Genomics) with default parameters and aligned to the Gallus gallus Ensembl reference genome (bGalGal1.mat.broiler.GRCg7b; GCA_016699485.1). Downstream analysis was performed in R using the Seurat package (v5). To enrich for virally labeled CNCCs, cells were retained only if the viral reporter H2B-RFP was detected (H2B-RFP counts > 0). To minimize erythrocyte contamination, cells with elevated hemoglobin expression were excluded by requiring that HBBA counts accounted for < 0.5% of total UMI counts per cell. Cells from all time points passing these filters were merged into a single Seurat object. Additional quality control retained cells with 5,000–10,000 detected genes. The filtered dataset was log-normalized (counts per 10,000 followed by log1p transformation). Highly variable genes were identified using the variance-stabilizing transformation (vst) method. Data were then scaled and subjected to principal component analysis (PCA), followed by graph-based nearest-neighbor detection, clustering, and UMAP dimensional reduction to assess sample composition and reporter distribution prior to downstream comparisons. Differential gene expression between experimental groups (HH24 LAL vs HH24 control) was performed using a Wilcoxon rank-sum test on log-normalized expression values implemented in Seurat. Genes were pre-filtered to those detected in at least 10% of cells in either group. Effect sizes were calculated as the difference in mean log1p-normalized expression between groups and converted to log2 fold change by dividing by log(2). P-values were adjusted using the Benjamini–Hochberg procedure. Genes were considered differentially expressed if they satisfied both FDR < 0.05 and |log2FC| ≥ 0.1.

### 4.6 CNCC Post-migratory Program Scoring

To quantify developmental progression of CNCCs, a post-migratory CNCC gene program score was calculated for each cell using the log-normalized RNA expression matrix (Seurat RNA assay, “data” layer). The post-migratory program was defined based on curated marker genes (HAND1, FOXC2, GATA6, TBX2). For each cell, the program score was computed as the mean log-normalized expression of genes in the marker set using sparse-matrix column means. Cells were annotated by developmental stage (HH24-HH26) and treatment group (LAL vs control) according to sample metadata. Differences in program scores between groups were visualized using violin plots with overlaid boxplots and evaluated using two-sided Wilcoxon rank-sum tests.

### 4.7 Protein–Protein Interaction Network and Functional Enrichment Analysis

To investigate functional relationships among differentially expressed genes, a subset of genes downregulated in HH24 LAL CNCCs was analyzed using the STRING database (v12.0). Fifty genes associated with cardiovascular development and neural crest differentiation were selected from the downregulated differentially expressed gene list and used to construct a protein–protein interaction (PPI) network using the Gallus gallus reference dataset. The resulting interaction network was exported and visualized in Cytoscape (v3.9). Network clustering was performed using the Markov clustering (MCL) algorithm to identify functionally related gene modules. Genes from each cluster were subjected to Gene Ontology (GO) enrichment analysis in STRING, and clusters were annotated according to the significantly enriched biological processes using a false discovery rate (FDR) threshold of 0.05. Functional enrichment analysis of the selected 50 genes set was performed using STRING Gene Ontology (GO) enrichment analysis. Enriched biological processes were identified using FDR < 0.05 following Benjamini– Hochberg correction. Enriched GO terms were visualized using STRING enrichment plots.

### 4.8 Immunohistochemistry and Image Analysis

After cryosectioning, embryo sections were washed in PBS to remove residual PFA. The tissue was permeabilized with 0.2% Triton X-100 in PBS, followed by incubation with primary antibodies diluted in PBST containing 1% BSA and 0.3 M glycine at 4 ^°^C overnight. Subsequently, the sections were washed three times with PBST (PBS with 0.1% Tween 20) and incubated with the appropriate secondary antibodies and DAPI diluted in the same buffer for 1 h at room temperature. The sections were washed three times with PBS, incubated with an autofluorescence quenching agent (TrueVIEW, vector laboratories), washed with PBS, dried, and mounted for fluorescent imaging. Imaging procedures were performed on the Echo Revolution Microscope with 4× (channel DAPI, FITC) and 10× objectives (channel DAPI, FITC, ATX RED) (***Table 1***). The images were analyzed using the ImageJ. Primary antibodies include: 1:70.4 Mouse monoclonal anti-bovine Troponin T, IgG2a (CT3) (AB528495, DSHB), 1:400 Mouse monoclonal anti-NH2 terminal synthetic decapeptide of alpha smooth muscle actin, IgG2a (A5228, Sigma), and 1:9.4 Mouse monoclonal anti-chicken Myosin Heavy Chain, IgG1 kappa light chain (ALD58) (AB528361, DSHB). Secondary antibodies include: Goat polyclonal anti-mouse IgG1 Alexa-568 (AB2535766, Molecular Probes) and Goat polyclonal anti-mouse IgG2a Alexa-568 (AB2535773, Molecular Probes).

**TABLE 1.** Microscope calibration and fluorescence filter sets used for imaging.

| Objective calibration |  |  |  |
| --- | --- | --- | --- |
| Objective | Pixel size (μm/pixel) |  |  |
| 4× | 0.892 |  |  |
| 10× | 0.357 |  |  |
| 20× | 0.178 |  |  |
| 40× | 0.089 |  |  |
| Fluorescence filter sets |  |  |  |
| Channel | Excitation (nm) | Emission (nm) | Dichroic (nm) |
| DAPI | 385/30 | 450/50 | 425 |
| FITC | 470/40 | 525/50 | 495 |
| ATX RED | 530/40 | 590/50 | 560 |

### 4.9 Lightsheet microscopy

A selection of LAL and control embryos underwent endo-DISCO light-sheet preparation as previously described (^64^).Briefly, the embryo was dissected away from the egg and transferred into warm Tyrode’s solution. Using micro-needles fashioned from pulled capillary tubes (0.75 mm ID) cut to 20-35 µm inner diameter via a microforge (Glassworx, St. Louis, MO) and a micromanipulator (model M3301L, World Precision Instruments) the embryo’s vascular system was flushed with warm Tyrode’s solution, endopainted ^65^ with FITC-poly-L-lysine and perfusion fixed with 4% (w/v) paraformaldehyde (PFA, Sigma-Aldrich, St Louis, MO, USA) to preserve inner vascular volumetric integrity. The embryo was subsequently dehydrated through a series of methanol steps and optically cleared in Ethyl cinnamate (ECi). Lightsheet imaging of the embryos was performed using a Zeiss Lightsheet Z.1 system (Carl Zeiss, Oberkochen, Germany). The samples were illuminated with a light-sheet thickness of 4-7 µm and imaged with a 5X objective. The in-plane resolution was kept between 1.3 and 2.6 µm, and the through-plane (z) resolution was kept between 1.9 and 3.5 µm.

## Abbreviations

OFT: outflow tract
CNCCs: cardiac neural crest cells
CHDs: congenital heart defects
PAAs: pharyngeal arch arteries.

## AUTHOR CONTRIBUTIONS

Conceptualization - SEL formal analysis - AF, RP, SEL Funding acquisition - SEL, Investigation - AF, RP, Metholody - AF,HM, SEL Visualization - AF, SEL, RP, HM Writing - AF, RP, HM, SEL

## ACKNOWLEDGMENTS

The authors thank Marianne Bronner, Ph.D., and her lab for supply of viral vectors and general support. We thank Andrew McCulloch, Ph.D. and Jennifer Stowe, Ph.D. for generous use of reagents and trainee support, Brian Aguado, Ph.D. and Zeinab Jahed, Ph.D. for use of equipment, UCSD Human Embryonic Stem Cell Core, and UCSD Institute for Genomic Medicine.

## SUPPORTING INFORMATION

Additional supporting information may be found in a separate document.

