## Supplemental Files for "Altered Cardiac Neural Crest Migration Patterning in a Left Atrial Ligation Model of Hypoplastic Left Heart Syndrome"

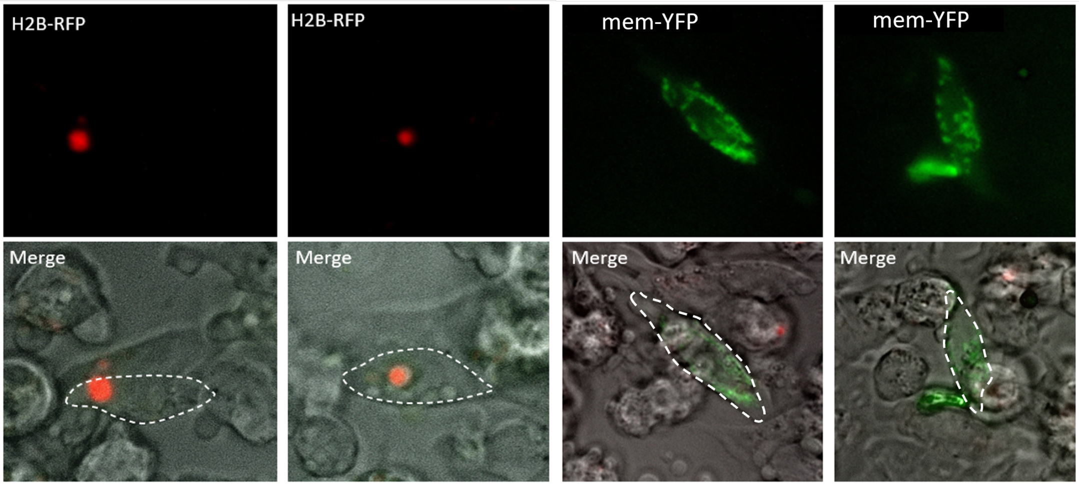


**Supplementary Figure 1: Validation of retroviral infection.** An embryo co-injected with RIA–H2B–RFP and RIA–membrane–YFP was dissected 48 h post injection. Cardiac tissues were enzymatically dissociated using Accumax at 37 °C to generate single-cell suspensions. Cells were collected in Hank's balanced salt solution supplemented with BSA, filtered through 40 $\mu$m cell strainers, and temporarily maintained in Hank's–BSA prior to imaging. Images were acquired using FITC and ATX-RED fluorescence channels together with transmitted light at 40× magnification. The dashed line outlines the cell membrane. H2B–RFP–transfected cells exhibit nuclear fluorescence, whereas membrane–YFP–transfected cells display fluorescence localized to the plasma membrane.

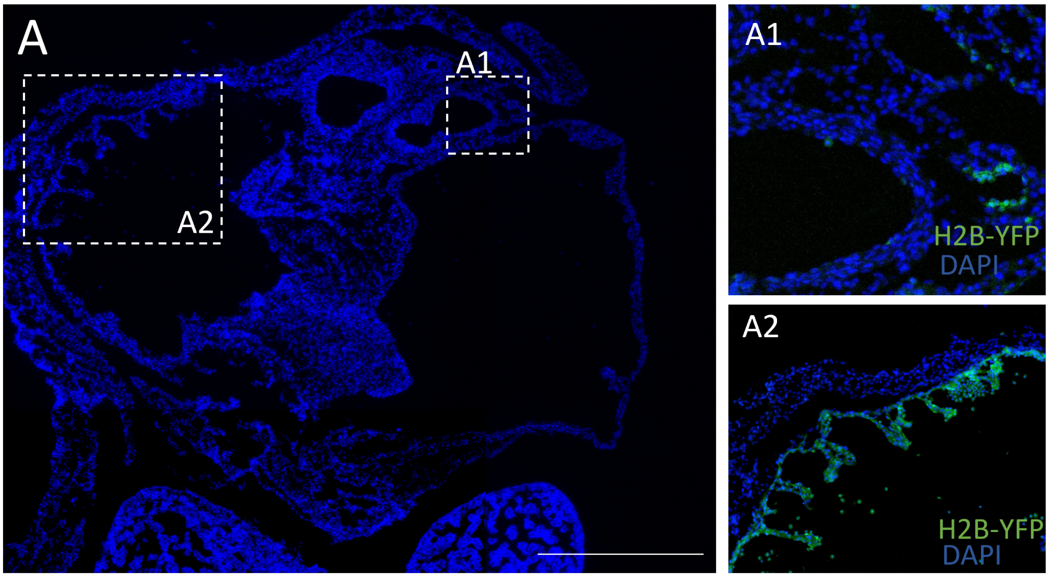


**Supplementary Figure 2: Distribution of CNCCs in ventricular myocardium and the OFT.** A single optical plane shows a region containing both ventricular myocardium and the OFT (***A***). Virally labeled CNCCs are detected in both structures(***A1***,***A2***). CNCC-derived cells are enriched within the ventricular myocardium, whereas little to no labeling is observed in the epicardial tissue surrounding the ventricle(***A2***). Scale bar: 500 $\mu$m

**TABLE 1** Quantification of CNCC distribution across tissue sections and anatomical regions. Sec1, Sec2, and Sec3 represent three consecutive sections separated by a z-stack interval of 10$\mu$m. CNCCs were identified and counted in each section, and mean values were calculated. The percentage of CNCCs in each anatomical region (PAA, OFT, MYO) was determined by dividing the number of CNCCs detected in that region by the total number of CNCCs identified across all three regions.

| | PAA | | | | OFT | | | | MYO | | | |
|---|---|---|---|---|---|---|---|---|---|---|---|---|
| Sample | sec1 | sec2 | sec3 | avg (%) | sec1 | sec2 | sec3 | avg (%) | sec1 | sec2 | sec3 | avg (%) |
| HH26 E1 | 169 | 140 | 158 | 155(81.5%) | 11 | 12 | 37 | 20(10.5%) | 8 | 18 | 21 | 15(7.8%) |
| HH26 E2 | 218 | 217 | 184 | 206(69.6%) | 52 | 75 | 89 | 72(24.3%) | 25 | 18 | 13 | 18(6.1%) |
| HH29 E1 | 324 | 198 | 209 | 243(44.5%) | 210 | 319 | 313 | 280(51.3%) | 18 | 39 | 16 | 22(4%) |
| HH29 E2 | 301 | 208 | 447 | 318(46.5%) | 258 | 256 | 543 | 352(51.5%) | 14 | 19 | 11 | 14(2%) |
| HH29 E3 | 309 | 270 | 220 | 266(67.9%) | 77 | 123 | 66 | 88(22.4%) | 20 | 67 | 29 | 38(9.7%) |
| LAL HH26 E1 | 90 | 112 | 146 | 116(57.1%) | 94 | 56 | 70 | 73(35.9%) | 29 | 6 | 9 | 14(6.8%) |
| LAL HH26 E2 | 182 | 166 | 171 | 173(75.9%) | 45 | 24 | 24 | 31(13.6%) | 26 | – | 22 | 24(10.5%) |
| LAL HH26 E3 | 81 | 91 | 106 | 92(66.2%) | 22 | 43 | 31 | 32(23.0%) | 22 | 13 | 12 | 15(1.1%) |
| LAL HH29 E1 | 127 | 146 | 166 | 146(78.5%) | 54 | 20 | 36 | 36(19.4%) | 2 | 3 | 9 | 4(2.2%) |
| LAL HH29 E2 | 158 | 114 | – | 136(76.8%) | 33 | 36 | 49 | 39(22.0%) | 1 | 4 | 3 | 2(1.1%) |
| LAL HH29 E3 | 150 | 228 | 176 | 184(74.2%) | 51 | 90 | 50 | 63(25.4%) | 1 | 1 | 0 | 1(0.4%) |

**TABLE 2** Selected differentially expressed genes (canonical CNCC migratory markers, HEY2, KLF2) identified in the HH24 Control vs HH24 LAL comparison. $Log_2FC$ represents the $\log_2$ fold change of gene expression in LAL relative to control. $pct_L$ and $pct_N$ indicate the fraction of cells expressing the gene in LAL and control samples, respectively.

| Gene | $\log_2$FC (LAL vs Control) | Adjusted p-value | $pct_L$ | $pct_N$ |
|---|---|---|---|---|
| SNAI1 | -0.886 | $2.57 \times 10^{-20}$ | 0.6738 | 0.8166 |
| HAND2 | -0.601 | $1.39 \times 10^{-5}$ | 0.4596 | 0.5976 |
| HEY2 | -0.505 | $1.41 \times 10^{-29}$ | 0.2482 | 0.6568 |
| ETS1 | -0.486 | $1.40 \times 10^{-14}$ | 0.6766 | 0.8402 |
| TFAP2A | -0.372 | $1.94 \times 10^{-6}$ | 0.8667 | 0.9586 |
| MAFB | -0.334 | $1.57 \times 10^{-10}$ | 0.3759 | 0.5917 |
| FOXC2 | -0.261 | $1.61 \times 10^{-12}$ | 0.3106 | 0.5976 |
| TAF10 | -0.260 | $5.05 \times 10^{-14}$ | 0.9801 | 0.9941 |
| HAND1 | -0.208 | $3.61 \times 10^{-10}$ | 0.1234 | 0.3195 |
| KLF2 | -0.105 | $2.18 \times 10^{-6}$ | 0.1050 | 0.2367 |
